# *In vitro* development and optimization of cell-laden injectable bioprinted gelatin methacryloyl (GelMA) microgels mineralized on the nanoscale

**DOI:** 10.1101/2023.10.10.560919

**Authors:** Mauricio Gonçalves da Costa Sousa, Gabriela de Souza Balbinot, Ramesh Subbiah, Rahul Madathiparambil Visalakshan, Anthony Tahayeri, Maria Elisa Lima Verde, Avathamsa Athirasala, Genevieve Romanowicz, Robert E. Guldberg, Luiz E. Bertassoni

## Abstract

Bone defects may occur in different sizes and shapes due to trauma, infections, and cancer resection. Autografts are still considered the primary treatment choice for bone regeneration. However, they are hard to source and often create donor-site morbidity. Injectable microgels have attracted much attention in tissue engineering and regenerative medicine due to their ability to replace inert implants with a minimally invasive delivery. Here, we developed novel cell-laden bioprinted gelatin methacrylate (GelMA) injectable microgels, with controllable shapes and sizes that can be controllably mineralized on the nanoscale, while stimulating the response of cells embedded within the matrix. The injectable microgels were mineralized using a calcium and phosphate-rich medium that resulted in nanoscale crystalline hydroxyapatite deposition and increased stiffness within the crosslinked matrix of bioprinted GelMA microparticles. Next, we studied the effect of mineralization in osteocytes, a key bone homeostasis regulator. Viability stains showed that osteocytes were maintained at 98% viability after mineralization with elevated expression of sclerostin in mineralized compared to non-mineralized microgels, indicating that mineralization effectively enhances osteocyte maturation. Based on our findings, bioprinted mineralized GelMA microgels appear to be an efficient material to approximate the bone microarchitecture and composition with desirable control of sample injectability and polymerization. These bone-like bioprinted mineralized biomaterials are exciting platforms for potential minimally invasive translational methods in bone regenerative therapies.

## 1. Introduction

Bone has a natural potential to self-regenerate. However, clinical defects larger than 2 cm are usually not self-repaired [1]. Therefore, repairing large bone defects remains a major challenge in regenerative engineering. Bone grafts remain the gold standard for treating traumatic bone injuries [2]. Nevertheless, autografts, allografts, and xenografts can cause infections, host rejections, and have limited availability [3]. Culturing osteoprogenitor cells on osteoconductive/osteoinductive scaffolds has been used as an alternative strategy for bone regeneration [4,5]. Despite the tremendous progress in cell therapy and bone tissue engineering, there is a limited number of biomaterials that can mimic the key characteristic defining the native bone microenvironment; that is, a cell-rich extracellular matrix where living cells are embedded in a heavily calcified matrix on the nanoscale [6,7]. Therefore, the goal of this in-vitro study was to determine and optimize the parameters that allow for engineering of a biomineralized and bioprinted bone-like biomaterial that may point to future translational applications.

Chief amongst the native composition of bone tissue, is the presence of a high concentration of osteocytes embedded in a collagenous scaffold [8]. It has previously been demonstrated that mineralization of such collagenous matrix follows a process typically referred to as non-classical mineralization, whereby negatively charged non-collagenous proteins (NCPs) interact with Ca^2+^ and PO_4_-3 [9] to enable the infiltration of ions into collagen fibrils, as an amorphous mineral phase, which then crystallizes intrafibrillarly as nanoscale needle-like particles [10]. This process results in the formation of what is known as the intrafibrillar collagen minerals, which has been shown to be responsible for most of the critical mechanical functions of bone tissue [11]. In previous work, we developed a method to mimic this process *in vitro* by rapidly mineralizing collagen hydrogels encapsulating mammalian cells of virtually any kind, to mimic the cellular and extracellular composition of native bone tissue *in-vitro* [12]. However, while this strategy mimics the composition of collagen in the bone matrix, it is limited by the associated difficulties of patterning microstructural hydrogels with fine geometrical control, or tunable physical properties, both which have been a major advantage of using light-based photopolymerization of photocrosslinkable hydrogels [13].

3D bioprinting of finely-resolved cell-laden photocrosslinkable hydrogels, in recent years, has benefited from a remarkable improvement in performance of bioprinted bone scaffolds enabled by digital light processing (DLP) bioprinting strategies [14]. These have ranged from biofabrication of photocurable bone-derived extracellular matrix, stackable LEGO-like bone grafts, multimaterial bioprinted hydrogels on a chip, only to name a few [15,16]. Moreover, these bioprinting strategies have been increasingly utilized to generate granular injectable hydrogels, which combine the benefits of minimally invasive cell-delivery, as well as the geometric control of biomaterial fabrication in high throughput [16].

While DLP bioprinting of photocrosslinkable hydrogels has been largely explored, including their use for bone regeneration [17], the ability of these biomaterials to mimic the cell-laden heavily calcified nature of bone tissue has remained virtually unaddressed. Here, we adapted our unique ability to calcify hydrogels loaded with living osteoprogenitor cells using a highly osteoinducitve approach, to generate injectable nanoscale mineralized cell-laden granular GelMA hydrogels bioprinted via DLP biofabrication. We demonstrate the fabrication and tunability of microgels bioprinted with different size and shape, as well as the biologic response of encapsulated cells in the bioprinted constructs. To further demonstrate the ability of the constructs to approximate the composition of bone autografts and allografts, both which are heavily populated by osteocytes at the time of delivery, we further studied the ability of these engineered constructs to be fabricated with dental pulp derived stem cells (DPSCs) and an osteocyte cell line (MLO-Y4). We hypothesized that fabrication of cell-laden bioprinted GelMA hydrogels followed by their nanoscale mineralization allowed for formation of heavily calcified injectable nanostructured hydrogels with matrix properties that are analogous to that of native bone tissue, inclusive of the presence of osteocytes embedded in minerals, and improved osteoprogenitor cell response *in-vitro*.

## 2. Experimental

### 2.1. GelMA synthesis

GelMA was synthesized using type A gelatin (10% w/v) from porcine skin (Sigma Aldrish, St. Louis, MO, USA) dissolved in PBS under stirring at 50°C. After gelatin dissolution, the methacrylic anhydride (Sigma Aldrish) was used at 8% (v/v) concentration dropwise for 2 h at 50 °C. The mixture was dialyzed against distilled water with 12–13 kDa tubes at 45 °C for five days with water changed twice a day. The GelMA solution was filtered with a 0.22 μm (Millipore Express Plus Stericups, Burlington, MA, USA) filter and stored at −80 °C overnight before lyophilization for five days.

### 2.2. Optimization of mineralization protocol

GelMA was dissolved in sterile phosphate buffered saline (PBS) at 10% (w/v) concentration for an initial screening for the ideal mineralization media. Printable hydrogels were produced with lithium phenyl (2,4,6-trimethyl benzoyl) phosphinate (LAP – L0290. Tokyo Chemical Industry, Tokyo, Japan) at 0.075% (w/v) concentration as initiator. Reagents were mixed for 60 s and maintained at 37 °C for 10 min for GelMA dissolution. Hydrogels (8 mm diameter x 1 mm thickness) were 3D printed in a stereolithography 3D printer (Ember. Autodesk, CA, USA) for 10 s under a 405 nm light at 20 mW cm^−2^ for 25 s. Immediately after printing, hydrogels were placed on 12-well plates for subsequent mineralization. Three different culture media were tested to optimize the cell medium composition used for nanoscale mineral formation: DMEM high glucose (4.5 g/l) (Thermo Fisher Scientific, Vancouver, BC, CA); DMEM low glucose (1g/l) (Thermo Fisher Scientific); and α-MEM (Thermo Fisher Scientific). All media were supplemented with 25 mM (4-(2-hydroxyethyl)-1-piperazineethanesulfonic acid) (HEPES, Sigma aldrish), 10% fetal bovine serum (Thermo Fisher Scientific), and 1% penicillin/streptomycin solution (Thermo Fisher Scientific). Mineralization was induced by subopplementing the media with 9 mM CaCl_2_. 2H_2_O (Fisher Scientific, Hampton, NH, USA) Osteopontin (100 mg/mL OPN. Lacprodan® OPN-10; Arla Foods Ingredients Group P/S, Denmark) as the NCP and 4.2 mM K_2_HPO_4_ (Fisher Scientific) as previously described [12]. The hydrogels were mineralized for three days with daily media changes. Rat-tail collagen (3 mg/mL, Thermo Fisher Scientific, Waltham, MA, USA) was mineralized under the same conditions as the GelMA hydrogels, as a positive mineralization control using our established methods [12]. Mineralization was assessed via Fourier-transform infrared spectroscopy (FTIR) to quantify the mineral matrix ratio, crystallinity index and to verify the presence of typical hydroxyapatite peaks. Spectra were obtained on a spectrometer (Nicolet 6700, Thermo Fisher Scientific) with 32 scans in a range between 400 and 4000 cm^−1^ in the transmission mode at 4 cm^−1^ resolution. The mineral matrix ration was calculated based on the band area of ν_3_PO_4_^3−^ (900– 1200 cm^−1^) and amide I (1660 cm^−1^), while the crystallinity index was calculated based on the doublet peak in the 500 and 650 cm^−1^ regions and the height of peaks and valleys on the ν_4_PO_4_^3−^.

### 2.3. Optimization of GelMA hydrogel parameters for improved mineralization

Using the optimized mineralization protocol, different GelMA parameters were investigated to assess differences in mineralization according to hydrogel concentration and cross-linking time. GelMA was dissolved in sterile PBS solution at 5, 7, 10 and 20% (w/v). Specimens (8 mm diameter x 1 mm thickness) were 3D printed under the abovementioned conditions for 25, 50, and 100 s. Hydrogels were characterized by their mineral matrix ratio and crystallinity index by FTIR (n = 3); and by their mechanical properties (n = 6), which was assessed under compressive load in a mechanical testing machine (Instron 5542 – Instron, Norwood, MA, USA) with a crosshead speed of 0.75 mm min^−1^. Stress and strain curves were used to calculate the elastic module based on the 5% initial strain in the linear region. Both mineralized and non-mineralized samples of GelMA as well as rat-tail collagen non-mineralized (3 mg/mL, BD Biosciences) were tested as a control.

### 2.4. Characterization of nanoscale mineralized GelMA

#### 2.4.1. Scanning electron microscopy

Mineralized GelMA (10% w/v) and collagen (3 mg/ml) specimens were fixed with 10% formalin solution for 10 min at room temperature and washed with distilled water. Samples were dehydrated in ascending ethanol series and critical point dried. A sputter-coating with palladium before imaging in an electron microscope (FEI Helios Nanolab™ 660 DualBeam™) at 15 kV. Elemental analysis was performed with an EDX detector (INCA, Oxford Instruments) for both mineralized and non-mineralized microgels.

#### 2.4.2. Transmission electron microscopy

Imaging was performed for GelMA (10% w/v) and collagen (1.5 mg/ml) hydrogels that were minced with a double-edge razor blade before immersion in ice-cold 0.1 M ammonium bicarbonate (pH 7.8). Minced hydrogels were then sliced in a tissue homogenizer (OMNI 2000, OMNI International, Kennesaw, GA) operated at ∼11,700 × g. Samples were pipetted onto freshly glow-discharged 600-mesh carbon-coated TEM grids and observed directly using FEI G20 TEM operated at 120 kV.

#### 2.4.3. Alizarin red

An alizarin red assay was performed to determine the presence of calcium phosphate deposition inside the microgels. The samples were stained with alizarin red solution 2% (w/v), incubated for 15 min, and washed three times. The stained areas were analyzed by inverted microscopy (FL Autos, EVOS – Thermo Fisher Scientific).

#### 2.4.5. Von Kossa staining

Von Kossa staining was performed to determine the presence of calcium or calcium salt in the microgels. For this purpose, microgels were washed with ultrapure water three times. The samples were incubated for 20 min in the UV light with 1% silver nitrate solution and washed with water 3x. The un-reacted silver was removed with 5% sodium thiosulfate for 5 min.

### 2.5. Printing specified geometries of GelMA for mineralization

To demonstrate the ability of bioprinting nanoscale mineralized GelMA hydrogels of different geometrical configurations, various shapes were fabricated and then subjected to mineralization. We designed and fabricated distinct micropatterns, ranging from squares, triangles, circles, hearts, stars, the OHSU logo, and a checkerboard pattern, using CAD (Fusion 360, Autodesk). CAD files were converted into image slices with the 3D printing software (print studio Autodesk). The overall width/diameter of the printed shapes ranged from 600 μm^2^ (squares) to 2.5 mm (OHSU and UO logo) and 450 μm in thickness. The shapes were printed under a light exposure of 50 s. The non-mineralized and mineralized microgels were then stained with a green dye (Procion MX Dye, Jacquard Products, Healdsburg, CA, USA) and alizarin red. The images were obtained by an inverted fluorescence microscope (FL Autos, EVOS-Thermo Fisher Scientific).

### 2.6. Cell culture and encapsulation

Human dental pulp stem cells (DPSCs, PT5025, Lonza, Bend, OR, USA) were cultured in α-MEM with 10% fetal bovine serum and 1% penicillin/streptomycin solution at 37 °C and 5% CO_2_. Murine osteocytes (MLO-Y4, Kerafast, Boston, MA, USA) were cultured α-MEM containing L-glutamine and deoxyribonucleosides, 5% fetal bovine serum, 5% calf serum, penicillin/streptomycin at 100 U/ml and 1% penicillin/streptomycin solution at 37°C, and 5% CO_2_ [18]. Microgels with GelMA 10% (w/v) were chosen for the following experiments.

#### 2.6.1 Cell viability

To test the toxicity of the bioprinted hydrogels, both DPSCs and MLO-Y4 cells were encapsulated in GelMA prepolymers at a concentration of 5.10^5^ cells/mL, the hydrogel was photocrosslinked for 50 s under a 405 nm light, and constructs were washed and incubated with live and dead cell assay die kit (Molecular Probes, Themo Fisher) for 10 min. Specimens were rinsed with PBS to remove the unreacted die and imaged in an inverted fluorescence microscope (n = 3) (FL Autos, EVOS – Thermo Fisher Scientific, Waltham, MA USA). Images were processed and analyzed by (ImageJ, FIJI) [19] to calculate the cell viability based on the ratio between the overall number of cells (blue/ 4’6’-diamino-2-phenylindole - DAPI stained) and dead cells (red/PI stained) after 3 (osteocytes) and 7 days (DPSCs).

#### 2.6.2 Cell morphology and function

Further studies were performed with MLO-Y4 cells, to characterize the ability of the mineralized constructs to enhance osteocyte-related features, since osteocytes represent the resident cells in the bone tissue that are actually entombed in a calcified matrix. The morphology of embedded osteocytes was assessed via actin (Molecular probes, Thermo Scientific) and DAPI through images taken in an inverted fluorescence microscope (FL Autos, EVOS-Thermo Fisher Scientific) after three days of mineralization. The number of cells, number of dendrites per cell and the length of dendrites per osteocytes in bioprinted microgels were quantified using Imaris 8.0.1 (Oxford Instruments, Tubney Woods, AB, UK) and ImageJ/FIJI [19] after 3 days of mineralization. The expression of the osteocyte-specific transcription factor, sclerostin (SOST), was assessed after 3 days of mineralization by confocal microscopy with a laser-scanning confocal microscope (Zeiss LSM 880, Oberkochen, Baden-Württemberg, Germany) for immunofluorescence imaging with samples fixed with 4% paraformaldehyde. Cells were permeabilized with 0.1% Triton X100, blocked with 1.5% bovine serum albumin in PBS, and incubated at room temperature for 1 h. Samples were then washed in PBS and incubated with the primary antibody (rabbit polyclonal SOST antibody (Novus Biologicals, NBP1-77461)) (1: 100 dilution), overnight at 4 °C. After washing with PBS, the samples were incubated with secondary antibody Alexa Fluor 555 goat anti-rabbit IgG (Thermo Fisher Scientific, A21244, 1: 500 dilution) for 2 h. The images were taken on a confocal microscope. The 3D reconstructions of z-stacks of samples were processed and rendered on ZEN Black (Zeiss) and Imaris 8 (Bitplane) software. The fluorescence intensity of sclerostin was quantified from a total of 9 individual images taken for three independent samples using ImageJ.

### 2.7. Microgels injectability

To verify microgels injectability through conventional syringe needles, 200 cell-laden microgels were suspended in 0.5mL of α-MEM with 10% fetal bovine serum and 1% penicillin/streptomycin solution with or without a green dye (Procion, MX Dye, Jacquard Products). Mineralized microgels were collected and injected in concavities created in polydimethylsiloxane (PDMS) to simulate an osseous defect (cylinder shape, 5 mm diameter) using a needle of 18G. The cellular viability was checked before and after the injection as described above.

### 2.8. Statistical analysis

Descriptive analysis was performed in SEM, TEM, and representative images from cell culture. Data were submitted to the Shapiro-Wilk normality test. One-way ANOVA and Tukey were used to analyze data from mineralization assays. Two-way ANOVA and Tukey were used to analyze elastic moduli, cell viability, cell morphology, and osteogenic induction for mineralized microgels. T test was used to compare cellular viability before and after injectability. All analyses were conducted under *α* = 0.05 by Graphad Prisma 6.0.

## 3. Results and discussion

Despite considerable progress in engineering biomaterials for bone regeneration, strategies that allow for biofabrication of hydrogels in a manner that replicates the native nanoscale calcification of the bone cell-laden mineralized ECM remains poorly explored. Here we developed a bioprinted cell-laden gelatin methacryloyl hydrogel that is mineralized on the nanoscale to promote robust osteocytic maturation of cells embedded in an engineered calcified matrix. We then sought to characterize the parameters that enable improved mineralization of cell-laden GelMA and the response of the embedded bone cells, which we propose will pave the way for future translational strategies on bone regeneration using osteocyte-embedded bone like biomaterials.

GelMA consists of gelatin that is functionalized with methacryloyl moieties, and is rich in small sequences of peptides, such as arginine-glycine-aspartic acid (RGD) and matrix metalloproteinase (MMP) binding sites [37], that facilitate cell adhesion and remodeling. Classically, collagen intrafibrillar mineralization happens in the presence of calcium, phosphate, and negatively charged non-collagenous proteins [23] binding to collagen. This interaction allows calcium and phosphate ions to penetrate the collagen fibrils by capillarity or electrostatic interactions, and then crystallize internally within the fibrils in the form of hydroxyapatite. On the other hand, extrafibrillar collagen mineralization happens when hydroxyapatite nucleates preferentially outside of to the collagen fibrils, in the presence or absence of nucleation inhibitors [38]. Our results demonstrated that, in the absence of collagen fibrils, the polymeric network of GelMA formed the templating structure for the penetration and coverage of minerals on the crosslinked structure of the hydrogel monomers. Based on these mechanisms of collagen intrafibrillar and extrafibrillar mineralization, the main hypothesis possibly explaining GelMA mineralization may be the nucleation and adsorption of supersaturated calcium and phosphate in solution that bind to gelatin polypeptides, either within or on the surface of the amorphous GelMA polymeric network. Therefore, to determine the parameters regulating the mineralization of cell-laden GelMA hydrogels, we first investigated the influence of media composition, GelMA concentration and crosslinking time on the fabrication of a bioprinted injectable bone-like cell-laden biomaterial.

### 3.1. GelMA mineralization

We first sought to optimize the parameters that controlled GelMA mineralization. We compared different printing times, GelMA concentrations, and mineralization media composition (Figure 1A). Among the screened media composition, the α-MEM-based mineralization media increased the mineral matrix ratio significantly compared to the low or high glucose DMEM media (Figure 1B, p<0.001), and crystallinity index compared to the low glucose group (Figure 1C, p<0.01). As described before, some components in α-MEM are not present in DMEM media, such as L-alanine, L-asparagine, L-aspartic acid, L-cysteine, L-glutamic acid, L-ascorbic acid, D-biotin, vitamin B12, Pyruvic acid, and Thioctic acid [20]. The presence of these amino acids and proteins in α-MEM but not in DMEM could explain the higher mineralization in the groups containing α-MEM [21].

**Figure 1.**
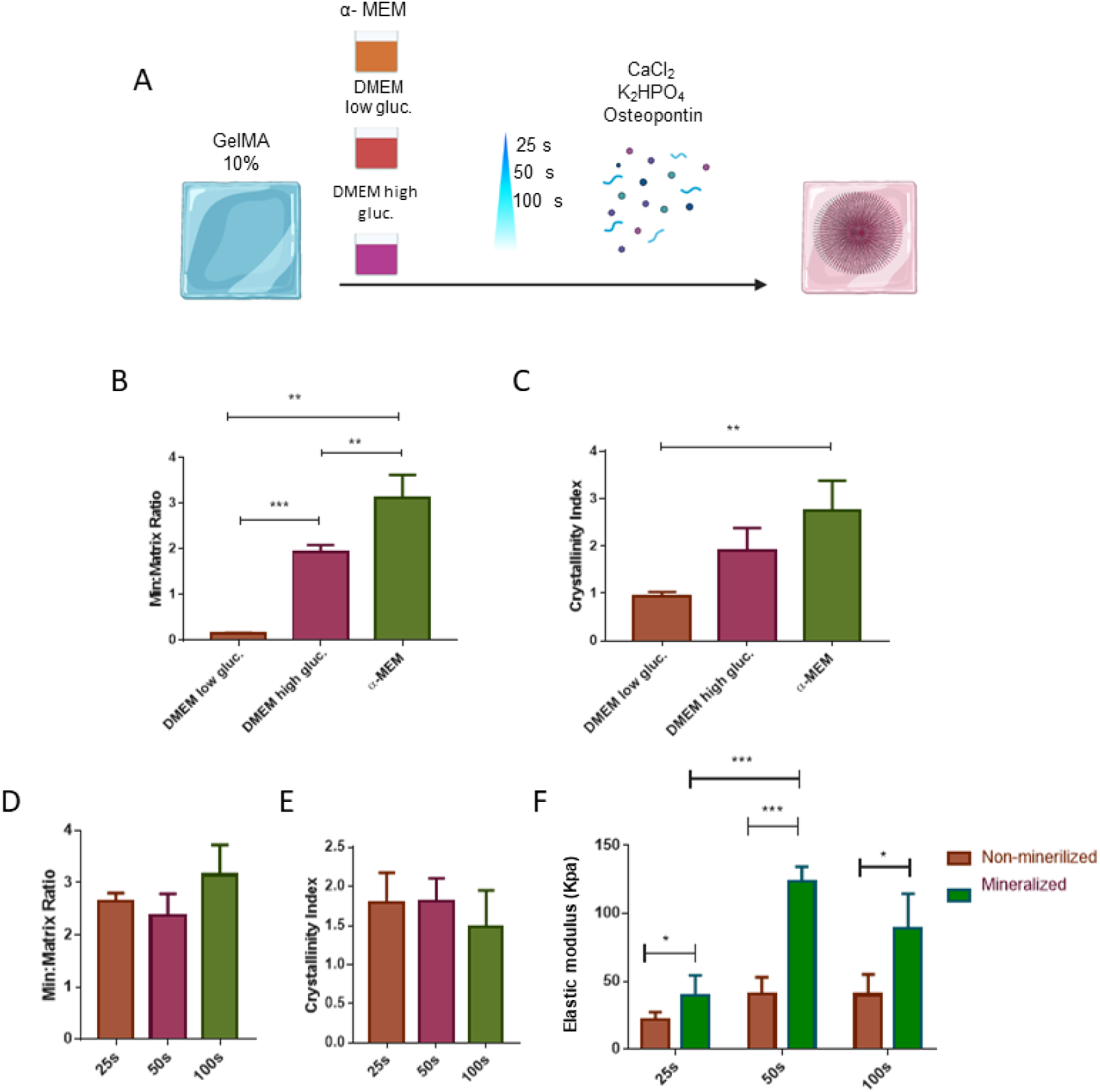
Optimization of different conditions for mineralizing microgels. (A) Microgels were mineralized with different conditions of photopolymerization time and media composition. (B) and (C) represent the mineralized matrix ratio and the crystallinity index of microgels cultivated in DMEM low and high glucose, as well as α-MEM media. (D-F) represent the mineralized matrix ratio, the crystallinity index, and the elastic moduli in kPa of microgels (GelMA 10%) that were exposed to 25, 50, or 100 seconds in the digital light printing (DLP) process. Statistical differences are represented by * (p<0.05), ** (p<0.01), and *** (p<0.001) after one-way or two-way ANOVA and a post-Tukey’s correction test.

Faced with α-MEM being the most effective media for mineralization, we tested different GelMA concentrations (5%, 7%, 10%, and 20%), and time points (25, 50, and 100 seconds) to understand how matrix concentration and crosslinking density would affect mineralization, as determined by mineral deposition and crystallinity index. We observed that polymerization time was a determinant factor in significantly modifying the stiffness of the groups containing GelMA 10 and 20% compared to the 5% and 10% groups (Figure 1D and supplementary figure 1). Based on these data, we choose GelMA 10% for our following experiments. As expected, in all time points (25, 50 and 100s), the elastic modulus was higher in the mineralized groups compared to the non-mineralized ones. However, when we compare just the mineralized groups, the GelMA stiffness reached a peak in 50 s (Figure 1). This finding might be explained by the saturation of GelMA crosslinking network, making less room from Ca^2+^ and PO_4_ biding.

### 3.2. Printability of GelMA microgels (tuning the size and shapes)

In order to demonstrate the patterning of microstructural hydrogels with fine geometry we printed the microgels with different shapes and sizes and mineralized those constructs. From our previous studies [22] and recent reports from other groups [23], it has been well demonstrated that the shape and size of granular microgels can affect cell response, osteogenic differentiation, and vasculature formation [24,25]. As a proof of concept of our ability to fabricate these geometric cell-laden microgels with nanostructural mineralization, we utilized CAD to design triangles, squares, circles, and the university (OHSU) logo with dimensions ranging from 600 µm − 2.5 mm. We then printed GelMA microgels using a DLP printer, as illustrated in Figure 2, and subjected them to the mineralization process described above.

**Figure 2.**
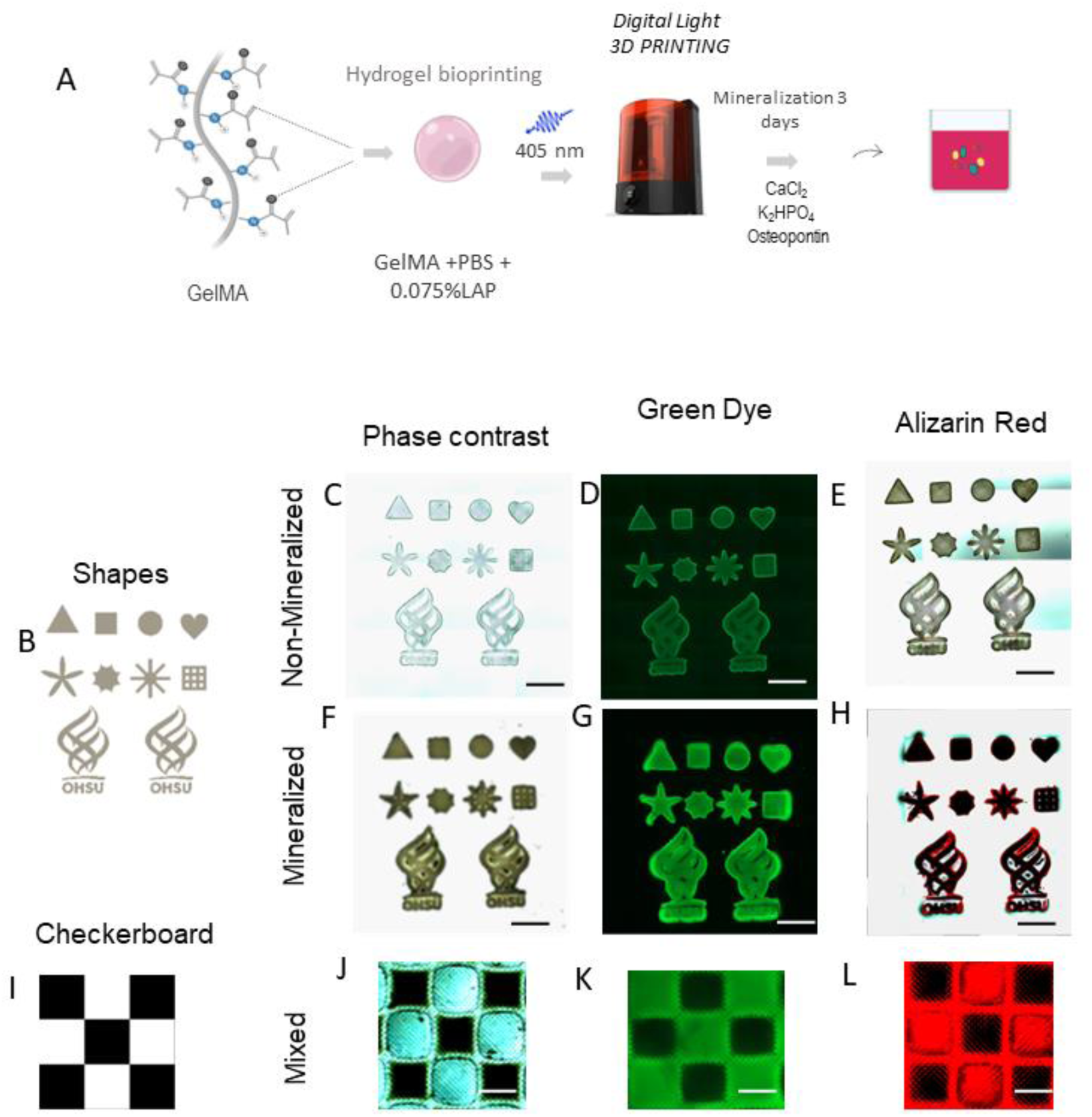
Printing and mineralization of microgels shapes. (A) Schematic of microgel preparation using DLP printer and mineralization steps. (B – H) represent the printed shapes, including triangle, square, circle, heart, stars, and the OHSU logo. (C, F, J) show the printed microgels in phase contrast. (D, G, K) show microgels imaged in green fluorescence. (E,H, L) show the printed constructs stained with alizarin red.

The microgels that were printed were stained either with green food color dye for easy visualization or with Alizarin red dye to confirm mineral deposition on the fabricated microparticles. We observed successful printing of these geometries as compared to the digital design, which were visualized using optical microscopy or the green food color dye with phase contrast or the green channel in the fluorescent microscope (Figure 2B-D). Alizarin red staining revealed that the mineralized constructs had dark red areas, indicating the effectiveness of our method in mineralizing different printed shapes throughout the fabricated areas (Figure 2F-H). Additionally, we demonstrated the ability to create heterogeneous constructs with both mineralized and non-mineralized regions in a checkerboard pattern (Figure 2I-L). We found that this pattern could be replicated by mineralizing microgels after three days and printing new microgels (non-mineralized) in empty spots. Optical microscopy of the green food dye confirmed the successful patterning of mineralized and non-mineralized microgels, resulting in a checkerboard panel (Figure 2J-K). Alizarin red staining confirmed the presence of calcium deposition in the mineralized zones of the checkerboard pattern (Figure 2L

### 3.3. Characterization of mineralized GelMA microgels

After printing microgels with different shapes and demonstrating that they could be mineralized, we selected square microgels for further characterization and biological activity. The mineralization via milk osteopontin (mOPN) in a supersaturated solution of Ca^2+^ and PO_4_ ^3−^ successfully promoted the infiltration of minerals into the GelMA structure, as observed in cross-sections of mineralized hydrogels in Figure 3 C and D by scanning electron microscopy, being distinct from non-mineralized samples (Figure 3C). When compared to collagen rat tail type I, mineralized GelMA structures were more disorganized (Supplementary Figure 2), showing that the mechanism of mineralization in GelMA might be different compared to collagen. Darker areas on TEM imaging denote the presence of hydroxyapatite crystals in the GelMA structure (Figure 3C).

**Figure 3.**
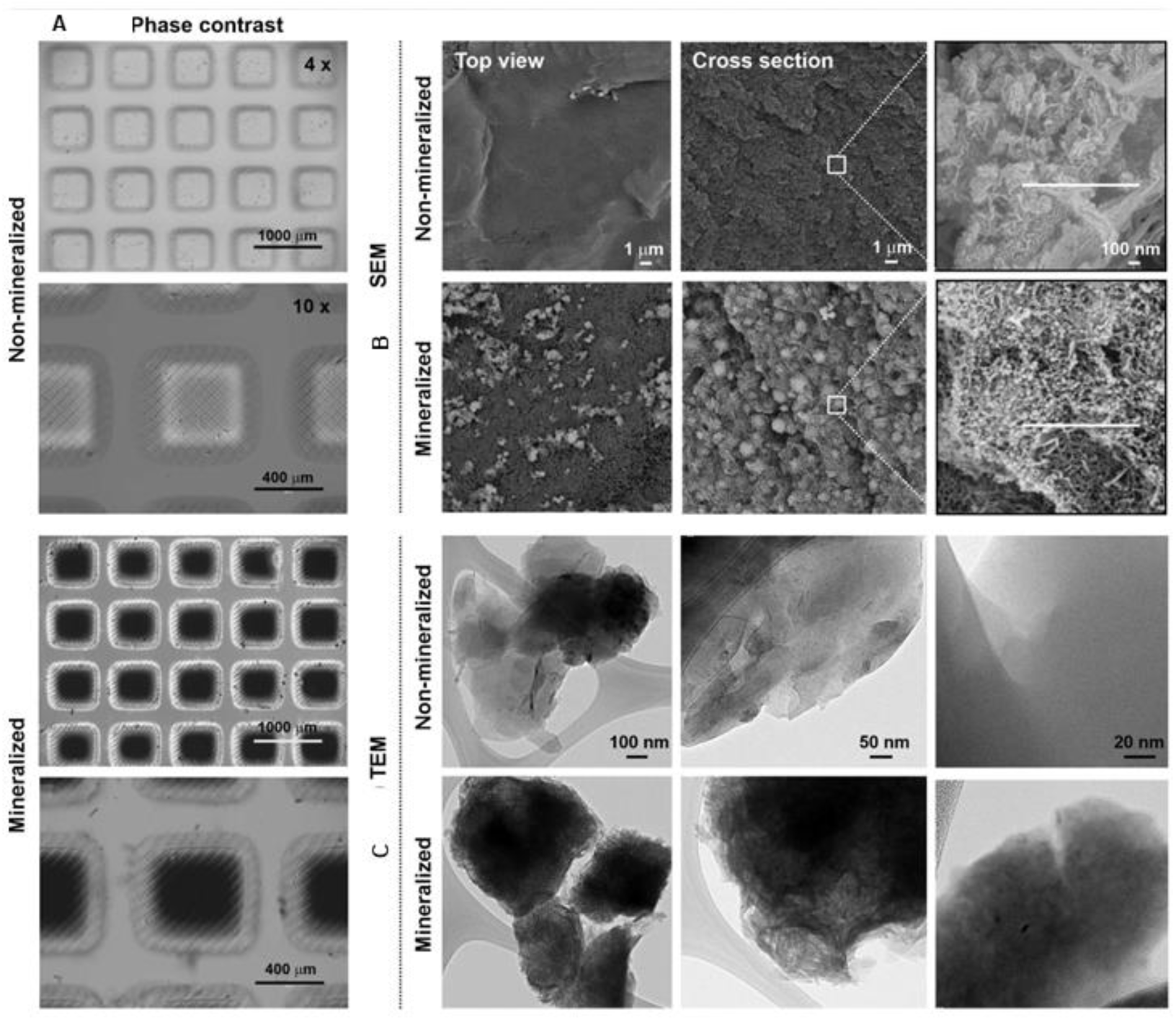
Microgel characterization. The images show the morphology of mineralized and non-mineralized microgels in 4 and 10x of magnification by light microscopy. B and C show the differences in morphology and extrafibrillar mineralization between mineralized and non-mineralized microgels from top view and from a cross-section view by scanning electron (B) and transmission electron microscopy (C).

Energy dispersive x-ray spectroscopy analysis demonstrated the presence of Ca and P only in mineralized microgels, confirming the successful mineralization of the samples (Figure 4C-D). Similarly, FTIR analysis of both mineralized and non-mineralized microgels demonstrated peaks consistent with amide I (1650 cm^-1^), amide II (1500 cm^-1^) and amide III (1240 cm^-1^) that are characteristic of the gelatin composition in GelMA (Figure 4D). However, just mineralized microgels demonstrated peaks absorbed between 500 to 700 cm^-1^ and 900 to 1020 cm^-1,^ consistent with phosphate apatite, confirming the presence of hydroxyapatite in the mineralized microgels (Figure 4D). Despite using an unique method to mineralize GelMA, our findings are consistent with other studies that characterized mineralized GelMA previously without cells in the matrix [27]. In summary, our data supports the conjecture that the mineralization after three days using CaCl_2_.2H_2_O, Osteopontin, and K_2_HPO_4_ approximates the process of mineral formation in calcified tissues like bone.

**Figure 4.**
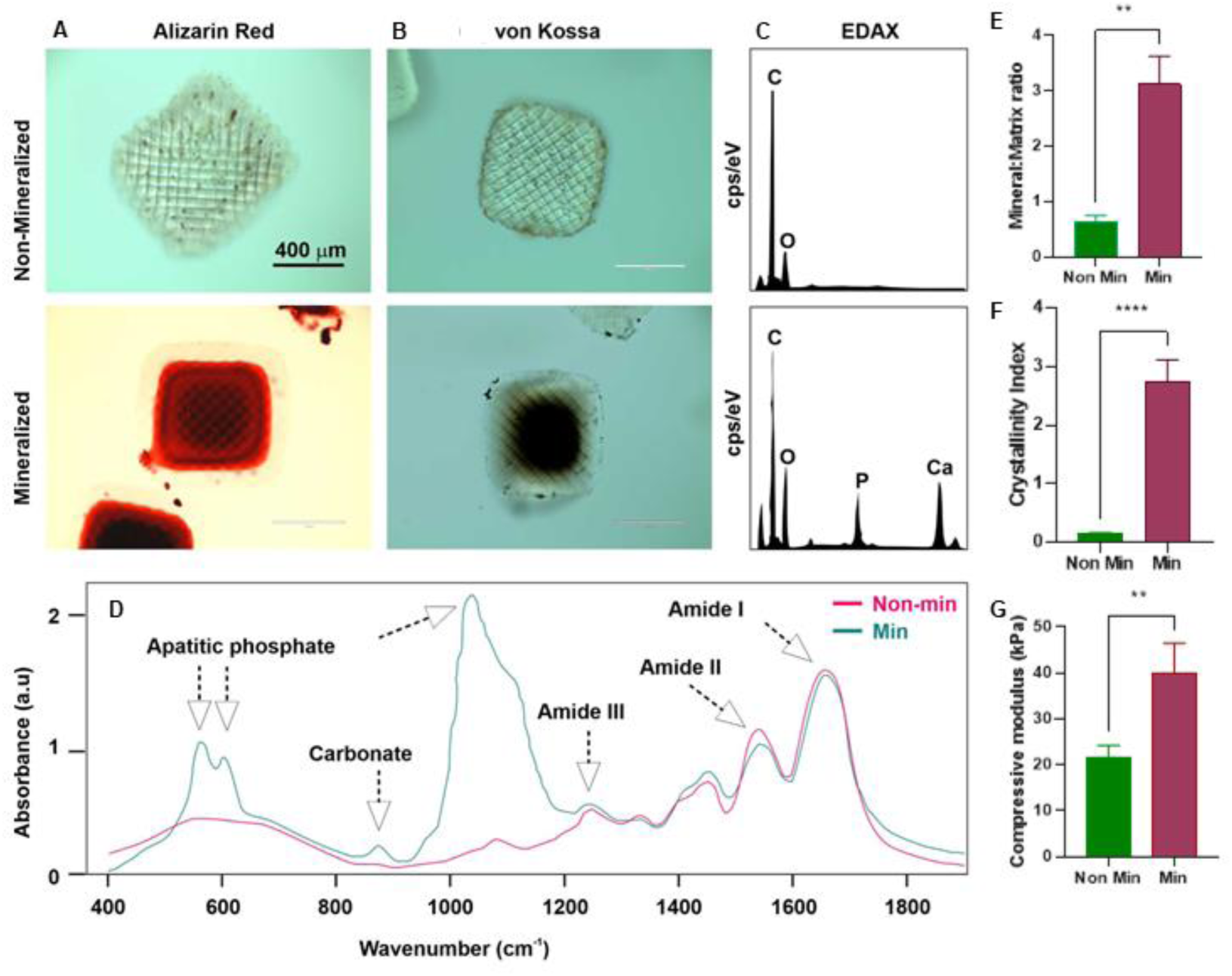
Characterization of non-mineralized and mineralized microgels. Mineralized microgels stained with Alizarin red (A) and von Kossa (B) show dark solid areas compared to the non-mineralized group after 3 days. By the energy diffraction x-ray analysis (EDAX), (C) the presence of Ca and P electron peaks were just observed in mineralized microgels. (D). FTIR analysis shows the absorbance of different organic (amides I-III) and inorganic (phosphate) chemical groups only on mineralized samples with the wavenumber expressed in cm-1. The mineral matrix ratio (E), the crystallinity index (F) and the elastic modulus confirm higher stiffness (represented in kPa) and solid composition of mineralized microgels compared to the non-mineralized samples. Statistical differences are represented by ** p<0.01, and **** p<0.0001 after the One-way ANOVA test post-Turkey’s corrections.

### 3.4. Biocompatibility and biological function of mineralized microgels

The bone microenvironment is characterized by different cells surrounded by a 3D calcified matrix where they biochemically interact with minerals, proteins, to perform their biological functions [28]. Hence, we replicated these bone-like mechanical cues in microgels, to engineer a proper microenvironment for bone cells. To that end, we evaluated the cellular behavior of the most abundant native bone cells (osteocytes) regarding their cell viability, proliferation, and maturation. We also evaluated the viability of more sensitive primary cells from the dental pulp (DPSCs) laden on mineralized microgels up to 7 days of incubation. Despite modifications in cell niche and the mechanical behavior promoted by minerals, mineralized hydrogels were able to maintain approximately 80% of DPSCs viable for at least three and seven days of incubation (Supplementary Figure 4). Additionally, 98% of the osteocytes were viable after three days of mineralization, proving that these mineralized structures did not cause cytotoxic effects for either cell type (Supplementary Figure 4).

We further explored the potential of these microgels to maintain osteocyte morphology and functionality after the mineralization process, since these cells are the major population in the bone calcified environment. For this purpose, we mineralized GelMA microgels for 3 days and stained these biomaterials with actin and DAPI. As expected, it was observed that the number of dendrites per cell and the dendrite length in the mineralized microgels were not significantly altered compared to the non-mineralized group (Figure 4 I-J). Interestingly, we observed that the number of osteocytes per microgel was approximately 3 times higher (p ˃ 0.001) in the mineralized microgels compared to the non-mineralized group, indicating that the mineralized environment stimulated osteocyte proliferation (Figure 4A, B, E, F, and K). We evaluated the expression of sclerostin (SOST) by immunostaining, a common protein produced by mature osteocytes. It was seen that the mineralized microgel group was responsible for stimulating higher SOST than the non-mineralized group (Figure 4G, H, L).

We chose the cell line MLO-Y4 since it is a well stablish cell line to study osteocyte behavior. These cells have been extensively reported in the literature as a model to study osteocyte physiology *in vitro* [18]. Our results demonstrate that these cells proliferated more in the mineralized microgels compared to the non-mineralized group. As a possible hypothesis for these findings, the stiffer matrix, where osteocytes are natively embedded has been demonstrated to influence osteocyte proliferation and function [29]. However, it is still controversial how the proliferation can be controlled in the 3D environment. According to Fournier 2020, MLO-Y4 cells cultivated in 3D had more cells in collagen droplets containing hydroxyapatite than just collagen after three days of culture [30]. It was also observed that the cell density could influence the dendritic profile in osteocytes. Collagen droplets with higher cell density (with HA) had a lower dendritic profile [30]. The proliferation of osteocytes in stiffer environments might be temporary once new findings show how that after 30 days of culture, the number of cells does not increase significantly. Yang, *et. al.,* 2020 bioprinted GelMA (5% w/v) and hyaluronic acid methacrylate (1% w/v) with IDG-SW3 osteocytes and evaluated the cell behavior after 30 days of culture. The authors observed that cellular viability was slightly reduced over time [31]. Sclerostin is a marker of mature osteocytes [32]. This protein is related to bone metabolism regulation, activating the Wnt/βcatenin signaling pathway [33]. Stiffness (mineralized matrix) has been demonstrated to influence the expression of SOST [34], and although this mechanism is not entirely understood, osteocytes have been proposed to have mechanosensors such as fibrils, integrin-based focal adhesion, ion channels, and connexin gap junctions, which could regulate sclerostin activation [32,35]. All of these mechanoregulatory factors point to an important participation of calcium, phosphate and their transformation into rigid minerals in the bone matrix as important determinants of osteocyte response.

### 3.6. Injectability

One of the significant challenges in the biomaterials field is developing products that are more easily translated into the clinic [38,39]. Among these barriers, injectability is still a challenge for bone tissue engineering substitutes [36]. Although some advances were achieved over time with injectable gels for bone tissue engineering, hydrogels that injected to conform to the defect sites in the form of monolithic materials do not provide the most favorable conditions for cell-cell interaction and nutrition [37,38]. Developing tiny bone-like cell-laden microgels might be an excellent opportunity to overcome these issues for hard tissue regeneration. In order to provide a platform to deliver cell-laden bone-like microgels, we tested the injectability of cell-laden microgels, and their ability to protect osteocytes during the shear stresses exerted on the material during injection. Cell-laden mineralized microgels were injected from an 18G needle to fill PDMS model-defects (Figure 5) that could artificially mimic a tissue cavity (1 mm in diameter). The cellular viability was evaluated before and after cell dispensing. After dispensing around 1 mL of microgels (resuspended in DMEM media), it was observed that the injectability process did not affect the osteocyte viability significantly (Figure 5G), showing that our biomaterial might be a good candidate for future translational applications, especially for bone defects (caused by traumas, infections, cancer, among other diseases) and bone augmentation (Figure 5H). Previously, another group have been demonstrated that the injection of cell laden microgels does not affect the viability of bone marrow cells (BMCs) after injectability into PDMS defects. The authors observed that just 15% of “naked” (not into microgels) BMCs injected with a needle in similar conditions were viable [39]. The fact that cells are embedded in a calcified environment might protect cells from shear stress during the injection, which could trigger cellular apoptosis [40], thus further suggesting that these novel bioprinted mineralized biomaterials hold promise for future therapeutic applications. While here we developed and characterized the parameters that enable the fabrication of injectable bone-like osteocyte-laden microgels, future work will focus on the use of this material for regenerative applications in in-vivo.

**Figure 5.**
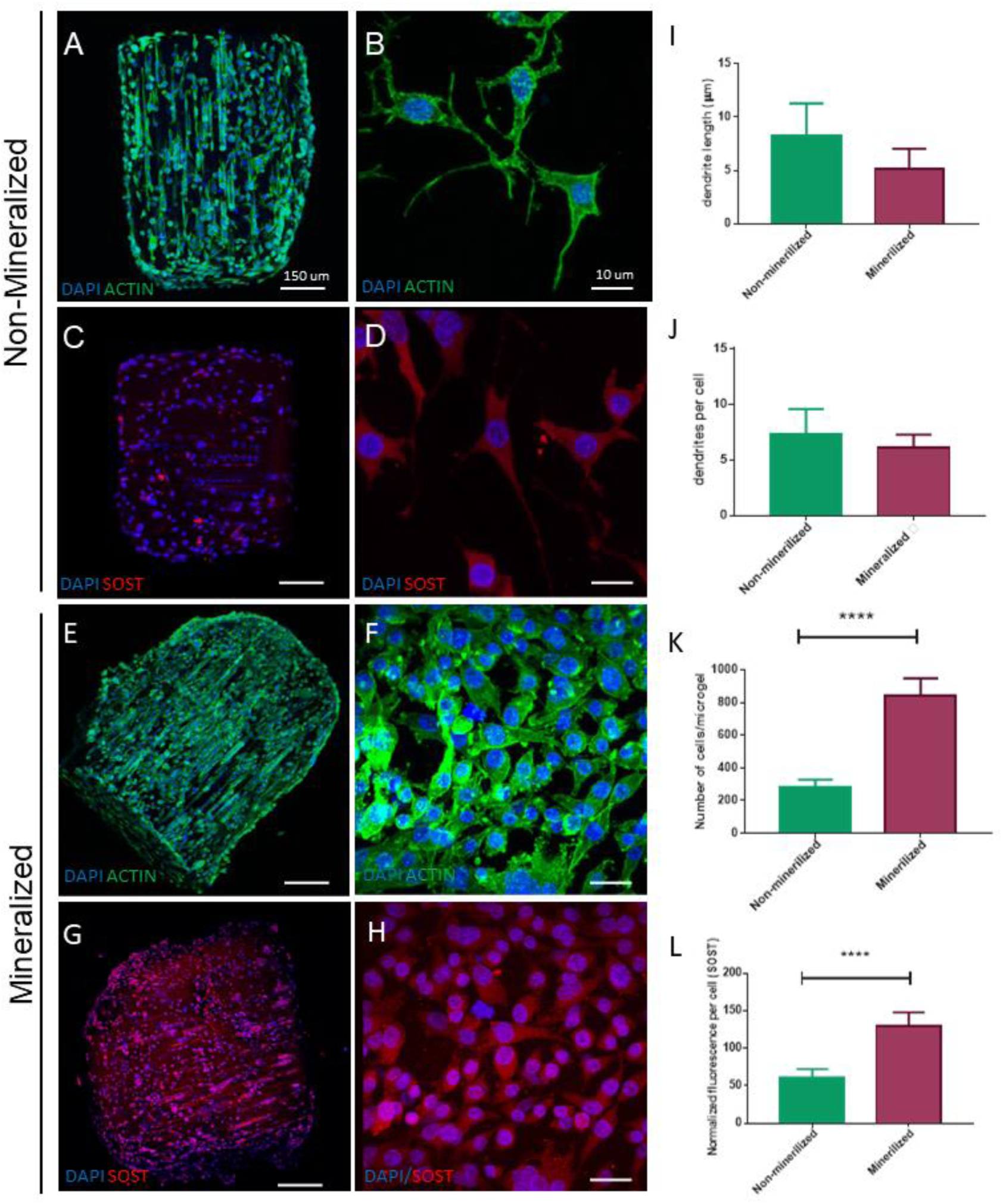
Osteocyte morphology and functionality in mineralized and non-mineralized microgels. A-B and E-F represent the confocal images of non-mineralized (A-B) and mineralized (E-F) samples. Microgels stained with actin, DAPI, and SOST revels higher cell number and SOST intensity (normalized fluorescence per cell) in mineralized microgels (A-H, K-L). Also, the dendrite length in μm (I) and the number of dendrites per cell (I-J) were not significantly different between mineralized and non-mineralized samples. Statistical differences are represented by **** p<0.0001 after the One-way ANOVA test post-Turkey’s corrections.

**Figure 6.**
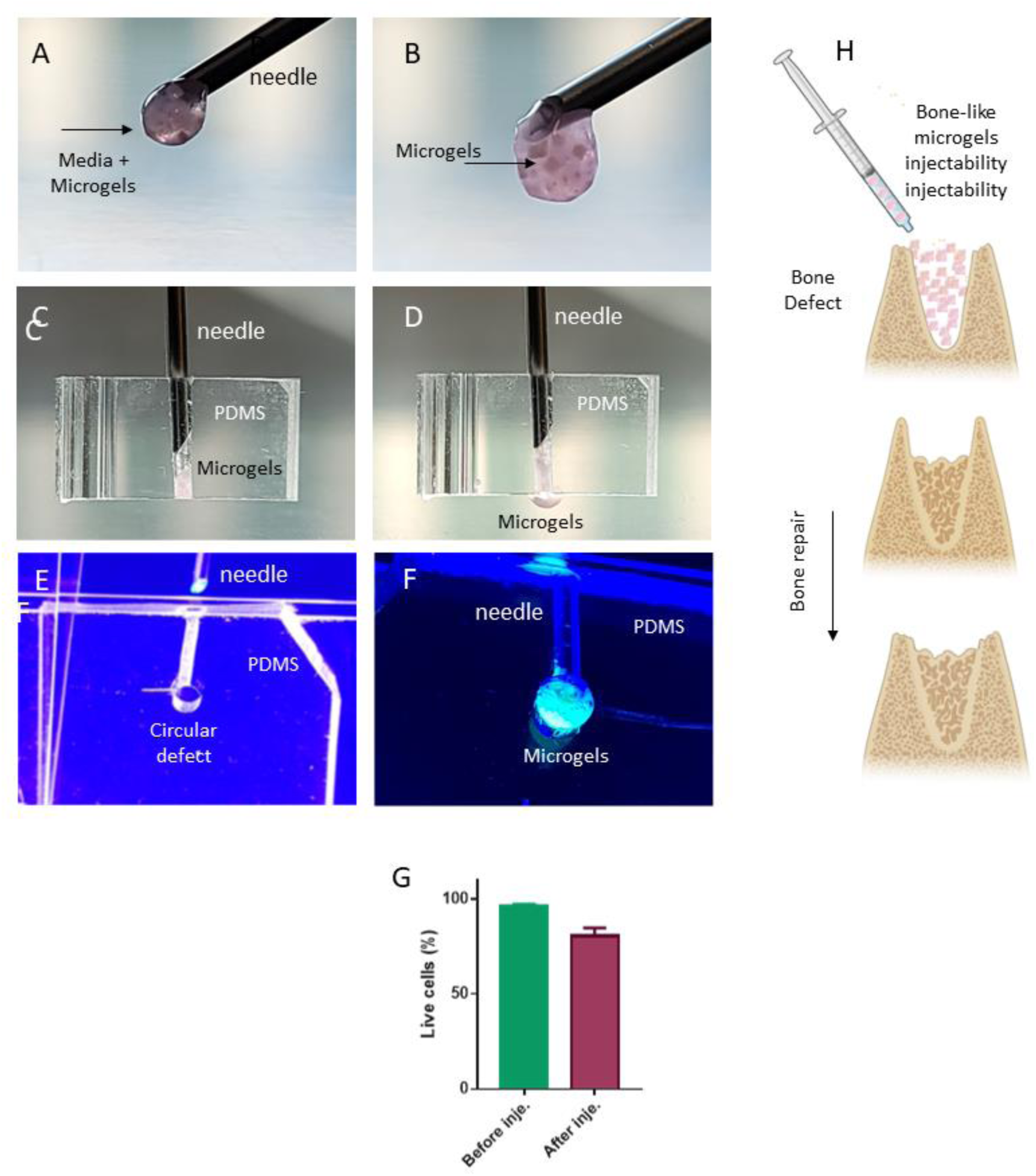
Microgels injectability. Microgels were mineralized for three days and injected from an 18 G needle (A-B) in PDMS defects (1mm of diameter) (C-F). The cellular viability of osteocytes in mineralized microgels was evaluated by the live and dead stain. The G Figure illustrates the percentage of live osteocytes on technical and biological triplicates before and after injection. There was no significant difference between the groups after t test analysis between the two groups.

These studies are currently ongoing and for the basis for future reports, which also points to a limitation of the present work.

## 4. Conclusion

In this study, we developed injectable nanoscale-mineralized bioprinted cell-laden GelMA microgels that demonstrated desirable maturation of an osteocyte cell-line and good translational potential for future therapeutic applications. Using several methods, we confirmed that the microgels were consistently mineralized, emulating the bone microenvironment composition. These bone-like biomaterials can be designed with controllable shape and stiffness, printed by digital light processing bioprinting with cells, and optimized for fine-needle injection, which could expand their applications for translational bone biofabrication.

## Acknowledgments

This project was supported by funding from the National Institute of Dental and Craniofacial Research (R01DE026170, 3R01DE026170-03S1, R01DE029553, and T90DE030859), Peter Geistlich Research Award (Osteo Science Foundation), and Oregon Clinical & Translational Research Institute (OCTRI) – Biomedical Innovation Program (BIP).

## Notes

### Competing Interest Statement

The authors have declared no competing interest.

