## Supplemental material for "*In vitro* development and optimization of cell-laden injectable bioprinted gelatin methacryloyl (GelMA) microgels mineralized on the nanoscale"

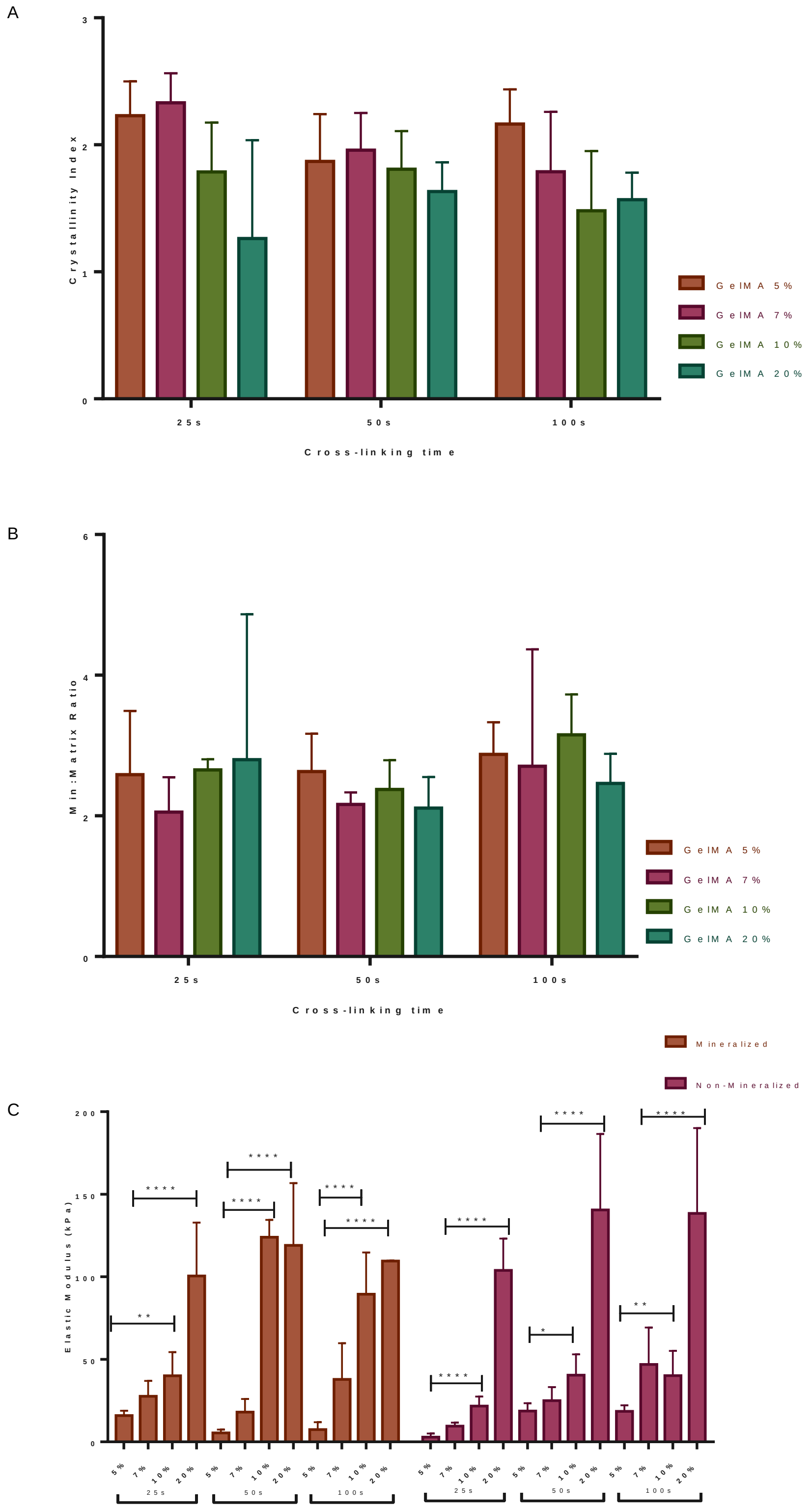


Supplementary figure 1 – Microgels optimization. The crystallinity index (A) and the mineral matrix ratio (B) were tested for mineralized GelMA microgels with different concentrations (5, 7, 10, and 20%) and time

(25, 50 or 100s). C represents the stiffness of these microgels with different concentrations and different concentrations (5, 7, 10, and 20%) and time (25, 50 or 100s) represented by the elastic module (Kpa). Statistical differences are represented by * (p<0.05), ** (p<0.01), *** (p<0.001), **** (p<0.0001) after two-way ANOVA test.

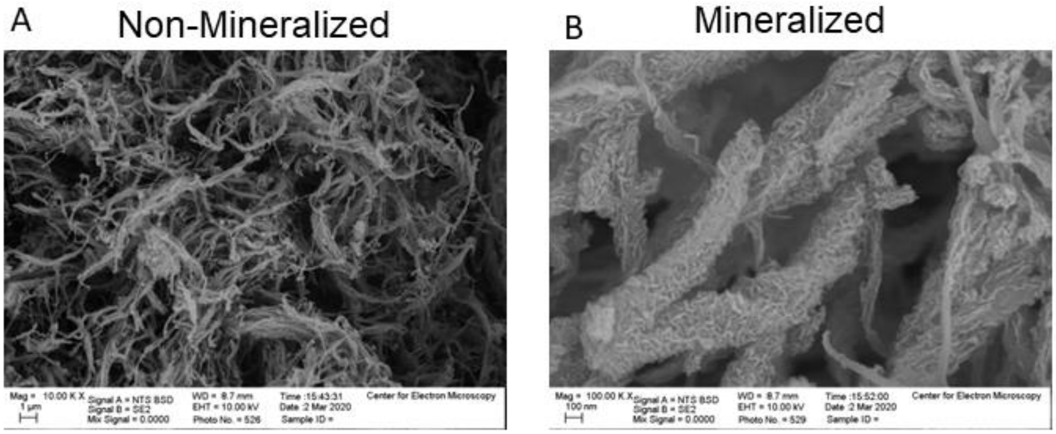


Supplementary figure 2 – Scanning electron microscopy of non-mineralized (A) and mineralized (B) collagen.

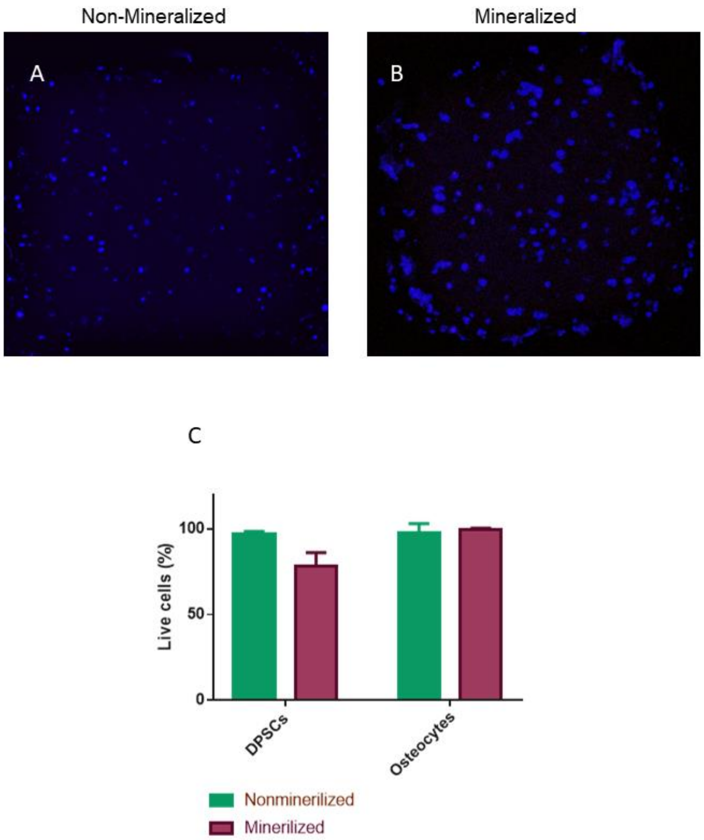


Supplementary figure 3 - Cellular viability in non-mineralized and mineralized microgels. Dental pulp stem cells (DPSCs) and osteocytes were cultivated for 3 days into mineralized and non-mineralized microgels. Bars represent the percentage of live cells of three different biological replicates.
